# A toxin-antidote CRISPR gene drive system for regional population modification

**DOI:** 10.1101/628354

**Authors:** Jackson Champer, Yoo Lim Lee, Emily Yang, Chen Liu, Andrew G. Clark, Philipp W. Messer

## Abstract

Engineered gene drives have been suggested as a mechanism for rapidly spreading genetic alterations through a population. One promising type of drive is the CRISPR homing drive, which has recently been demonstrated in several organisms. However, such drives face a major obstacle in the form of resistance against the drive that typically evolves rapidly. In addition, homing-type drives are generally self-sustaining, meaning that a drive would likely spread to all individuals of a species even when introduced at low frequency in a single location. Here, we develop a new form of CRISPR gene drive, the Toxin-Antidote Recessive Embryo (TARE) drive, which successfully limits resistance by targeting a recessive lethal gene while providing a recoded sequence to rescue only drive-carrying individuals. Our computational modeling shows that such a drive will have threshold-dependent dynamics, spreading only when introduced above a frequency threshold that depends on the fitness cost of the drive. We demonstrate such a drive in *Drosophila* with 88-95% transmission to the progeny of female drive heterozygotes. This drive was able to spread through a large cage population in just six generations following introduction at 24% frequency without any apparent evolution of resistance. Our results suggest that TARE drives constitute promising candidates for the development of effective, regionally confined population modification drives.

## INTRODUCTION

Gene drives have the potential to rapidly spread through a population by biasing inheritance in favor of the drive allele^1–7^. These systems could be used for population modification by carrying a payload allele engineered for a particular purpose, such as a transgene that reduces the capacity for malaria transmission in mosquitoes^1–3,5^. Gene drives may also be used for the direct suppression of a population, for example by targeting an essential gene of recessive effect. Such suppression-type drives could potentially be deployed against disease vector populations, invasive species, or agricultural pests^1–3,5^.

CRISPR-based homing drives promise a flexible gene drive mechanism for both population modification and suppression, and such systems have now been demonstrated in a variety of organisms, including yeast^8–11^, flies^12–18^, mosquitoes^19–21^, and mice^22^. These constructs work by cleaving a wild-type allele at a predetermined target site. The drive allele is then copied into the cleaved site during homology-directed repair, converting heterozygotes for the drive allele into homozygotes in their germline. However, the spread of CRISPR homing drives is typically thwarted by the formation of resistance alleles when Cas9 cleavage is repaired by end-joining, which tends to generate indel mutations at the target site^15^. This can take place both in the germline as an alternative to drive conversion and during early embryo development due to cleavage by maternally-deposited Cas9^15^. While several strategies for reducing the rate of resistance allele formation have already been successfully tested, including gRNA multiplexing^16^ and improved promoters^16,23^, these improvements have not been sufficient to reduce resistance rates to an acceptably low level.

Recently, a CRISPR-based population suppression drive that combined an improved promoter with a carefully selected target site where resistance alleles are non-viable was shown to be successful at suppressing a small cage population of *Anopheles gambiae*^24^. While promising, such a strategy may not be easily adoptable for approaches where the aim is population modification rather than suppression, and computational modeling has indicated that even high-performance population suppression systems may still face substantial evolutionary and ecological obstacles^25^. CRISPR homing gene drives also require Cas9 cleavage in the germline during a narrow temporal window, allowing for homology-directed repair instead of end-joining, as the latter will typically lead to the formation of resistance alleles. This increases development difficulty when designing homing drives in new species due to the need for a suitable promoter. Thus, new types of flexible population modification systems that minimize formation of resistance alleles may be needed for use either alone or in combination with a population suppression system.

One possible strategy for reducing resistance potential is to remove the need for homology-directed repair altogether. This criterion is fulfilled by drives using the “toxin-antidote” principle. Natural examples of such systems are numerous^26^, and a toxin-antidote system named *Medea* was successful in *Drosophila melanogaster*^27^. However, because *Medea* uses elements that are specific to *Drosophila*, it has proven difficult to move to other species. Other designs for engineered toxin-antidote systems also exist^28–33^, but elements for construction of such systems would likely be difficult to identify. One possibility for how such a system could be constructed is by engineering a drive allele that contains Cas9 and gRNAs serving as the “toxin” by targeting a recessive lethal (or strongly deleterious) gene. The drive allele would also contain an “antidote”, consisting of a recoded copy of the target gene that avoids homology to the gRNA. Such a drive would steadily convert wild-type target alleles in a population to disrupted alleles, at which point they would be removed from the population in embryos where no drive or wild-type allele is present to provide rescue. We term such a system TARE (Toxin-Antidote Recessive Embryo) drive.

In addition to minimizing resistance, a TARE system would also be expected to exhibit threshold-dependent invasion dynamics. This is in contrast to homing-type gene drive systems, which provide little control over the spread of a drive once released due to their ability to invade any distant regions with even a small number of migrants. Homing drives are therefore considered “global” drives^34^. This is often undesirable for various reasons, for example in conservation applications where the drive needs to be confined to a specific island^35^. TARE drives would likely remain regionally confined to contiguous population areas, without being able to establish in distant populations through occasional migrants.

A recent study has provided the first demonstration of a “distant-site” TARE drive, termed ClvR, where the drive allele is not at the same locus as its target site^36^. ClvR was able to successfully spread through small population cages^36^. Here, we demonstrate a “same-site” TARE system, which may have several advantages over the distant-site system, including efficiency of rescue, since the rescue element is at the same genomic site and uses the target gene’s natural promoter elements. We show that our TARE system successfully biases inheritance of the drive allele and reaches all individuals in a large *Drosophila* population cage after just six generations following a modest size release, without any apparent formation of resistance alleles.

## METHODS

### Simulations

Deterministic, discrete-generation simulations were initialized by seeding a population of wild-type individuals with drive/wild-type heterozygous individuals at a specified introduction frequency. Each female individual selects a mate randomly in each generation, where the probability of a male to be chosen is proportional to its fitness value. Females then generate a number of offspring equal to twice their fitness value. Fitness costs per drive allele are assumed to be codominant and multiplicative. In this model, fitness values thus represent fecundity for females and mating success for males relative to wild-type individuals. For the TARE system and the homing drive, we model both germline and embryo cleavage events. Disruption of the target gene by the TARE system occurs in both the germline and the embryo at 100% efficiency. Drive conversion for the homing drive occurs only in the germline of drive/wild-type heterozygotes at 100% efficiency. All offspring with non-viable genotypes due to TARE or *Medea* effects are then removed. Offspring genotype frequencies are then renormalized to generate the final allele frequencies at the end of the generation.

### Plasmid construction

The starting plasmids pCFD3^37^ (Addgene plasmid #49410) and pCFD5^38^ (Addgene plasmid #73914) were kindly supplied by Simon Bullock, and starting plasmid IHDyi2 was constructed in our previous study^15^. All plasmids were digested with restriction enzymes from New England Biolabs (HF versions, when possible). PCR was conducted with Q5 Hot Start DNA Polymerase (New England Biolabs) with DNA oligos and gBlocks from Integrated DNA Technologies. Gibson assembly of plasmids was conducted with Assembly Master Mix (New England Biolabs), and plasmids were transformed into JM109 competent cells (Zymo Research). Plasmids used for injection into eggs were purified using the ZymoPure Midiprep kit (Zymo Research). Cas9 gRNA target sequences were identified using CRISPR Optimal Target Finder^39^. Tables of DNA fragments used for Gibson Assembly of each plasmid, PCR products with the oligonucleotide primer pair used, and plasmid restriction digests with the restriction enzymes are shown in the Supplemental Information.

### Generation of transgenic lines

Lines were transformed at Rainbow Transgenic Flies by injecting the donor plasmid (EGDh2) into a *w*^*1118*^ line. Plasmid pHsp70-Cas9^40^ (provided by Melissa Harrison & Kate O’Connor-Giles & Jill Wildonger, Addgene plasmid #45945) was included as a source of Cas9 and plasmid EGDhg2t was included as a source of gRNA in the injection. Injection concentrations of donor, Cas9, and gRNA plasmids were 314, 313, and 63 ng/µL, respectively in 10 mM Tris-HCl, 100 µM EDTA, pH 8.5 solution. To obtain homozygous lines, injected individuals were crossed with *w*^*1118*^ flies. The progeny with dsRed fluorescent protein in the eyes, which usually indicated successful insertion of the drive, were crossed with each other for several generations, with preference to flies with slightly brighter eyes, which usually indicated that the individual was homozygous for the drive. The stock was considered homozygous after sequencing confirmation. The split-CRISPR line with Cas9 driven by the *nanos* promoter was generated as part of a previous study^17^.

### Fly rearing and phenotyping

All flies were reared at 25°C with a 14/10 hr day/night cycle. Bloomington Standard medium was provided as food every two weeks. For phenotyping, flies were anesthetized with CO_2_ and examined with a stereo dissecting microscope. Red fluorescent eye phenotypes were scored using the NIGHTSEA system (SFA-GR). The different phenotypes and genotypes of our drive system are summarized in Datasets S1-S2, as are the calculations we used for determining drive performance parameters.

For the cage study, enclosures of internal dimensions 30×30×30 cm (Bugdorm, BD43030D) were used to house flies. At the start of an experiment, drive flies and split-Cas9 flies were crossed as above until found to be homozygous at both sites by higher red and green fluorescent brightness and confirmed by subsequent crosses with wild-type individuals. These, together with split-Cas9 flies of the same age, were separately allowed to lay eggs in eight food bottles for one day. Bottles were then placed in cages at the desired starting ratios between drive and non-drive flies. Eleven days later, bottles were replaced in the cage with fresh food, leaving adult flies in the cages. One day later, bottles were removed from the cages, the flies were frozen for later phenotyping, and bottles were returned to the cage. This 12-day cycle was repeated for each subsequent generation.

All experiments involving live gene drive flies were carried out using Arthropod Containment Level 2 protocols at the Sarkaria Arthropod Research Laboratory at Cornell University, a quarantine facility constructed to comply with containment standards developed by USDA APHIS. Additional safety protocols regarding insect handling approved by the Institutional Biosafety Committee at Cornell University were strictly obeyed throughout the study, further minimizing the risk of accidental release of transgenic flies. All drive flies also utilized our split-Cas9 system^17^, which should prevent the spread of the drive in the case of an accidental escape. Finally, the drive has threshold-dependent kinetics by design, which means that even a non-split version of the drive should not be able to spread in a population when released at low numbers.

### Genotyping

To obtain the DNA sequences of gRNA target sites, flies were frozen and then homogenized in 30 µL of 10 mM Tris-HCl pH 8, 1mM EDTA, 25 mM NaCl, and 200 µg/mL recombinant proteinase K (Thermo Scientific). The mixture was incubated at 37°C for 30 min and then 95°C for 5 min. The solution was used as the template for PCR to amplify the gRNA target site. DNA was purified by gel extraction and Sanger sequenced. Sequences were analyzed using the ApE software, available at: http://biologylabs.utah.edu/jorgensen/wayned/ape.

## RESULTS

### TARE drive mechanism

The “same-site” TARE (Toxin-Antidote Recessive Embryo) drive consists of a drive element placed inside a recessive lethal gene. Though the presence of the drive disrupts the wild-type version of the gene, the drive construct contains a recoded version of a portion of this gene sufficient to restore its function, as well as a set of gRNAs that target only the wild-type gene at one or more target sites. Cleavage of the target gene by the drive creates a disrupted allele (typically termed “r2 resistance allele” in studies on homing drives, which are distinguished from the “r1” resistance alleles that maintain gene function). Individuals that possess two such disrupted alleles will be non-viable, leading to the systematic removal of such alleles from the population (Figure 1). As a result, the relative population frequency of the drive allele over the wild-type allele will increase over time.

**Figure 1.**
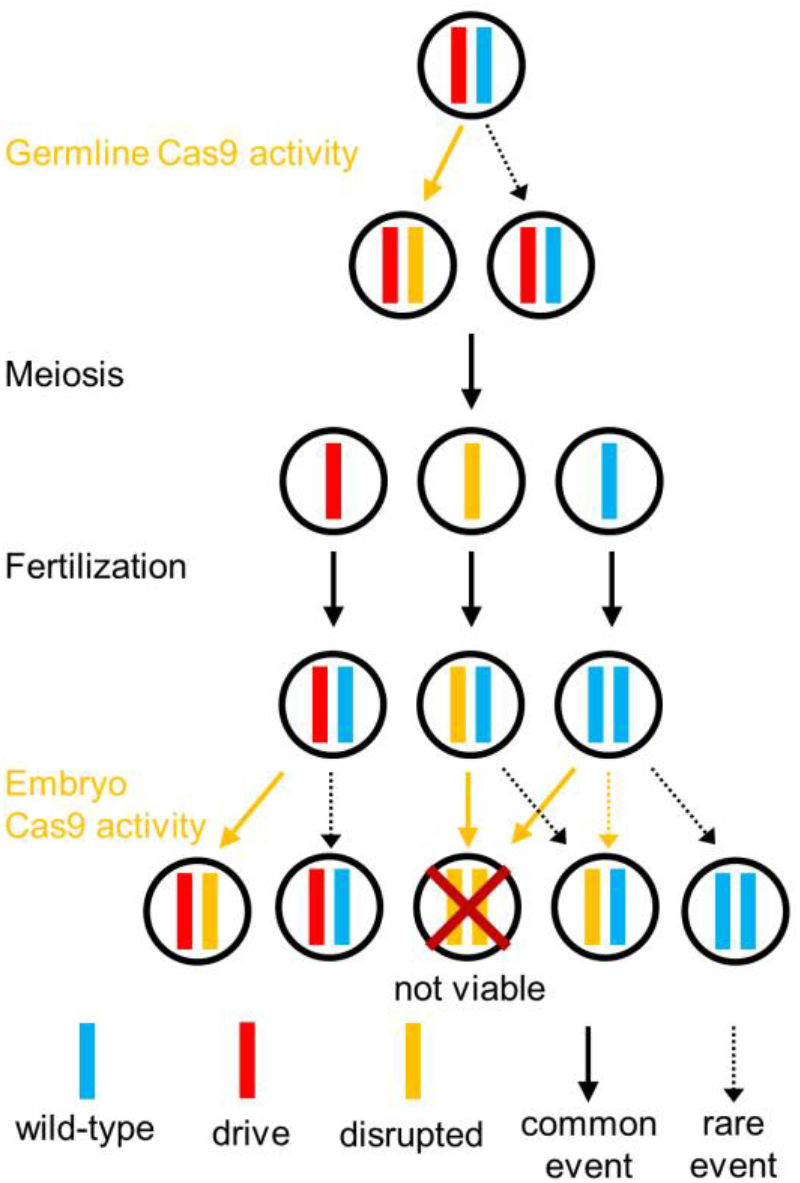
Mechanism of the TARE drive. In the germline of drive/wild-type heterozygotes, wild-type copies will usually undergo cleavage followed by homology-directed repair or end-joining, either of which will result in a disrupted target gene and loss of function. Meiosis and fertilization (shown here by a wild-type individual) then occur. In the progeny of females with the drive, maternally deposited Cas9 and gRNA will cleave most wild-type alleles, which will become disrupted after end-joining repair. Any individual that inherited two disrupted copies of the recessive lethal gene will be non-viable, which will lead to a systematic increase of the relative frequency of the drive allele over time.

The target site of the TARE drive needs to be sufficiently different from the one at which the drive is introduced to prevent the drive from being copied into the wild-type gene by homology-directed repair after cleavage. The recoded portion of the drive must further be designed such that it cannot be targeted by the drive gRNAs, nor have sequence homology around these cut sites. When cleavage of the target gene occurs in the germline, this can create a disrupted gene either through end-joining repair or by homology-direct repair around the cut site in the drive chromosome, converting the wild-type allele to the disrupted sequence form used for the template. In the progeny of females with the drive, maternally deposited Cas9 and gRNA will also result in cleavage of wild-type alleles in the embryo, creating disrupted alleles by end-joining and perhaps occasionally homology-directed repair^24^. Note that such a drive may additionally contain a payload gene.

### TARE drive population dynamics

Computational modeling suggests that a TARE drive will generally spread more slowly than a homing drive and instead have dynamics similar to a *Medea* system, although spreading somewhat more quickly (Figure 2A). All individuals rapidly become drive carriers with at least one copy of the TARE drive, particularly at higher release frequencies, but it can take quite long for the drive to eventually reach its maximum allele frequency (Figure 2B).

**Figure 2.**
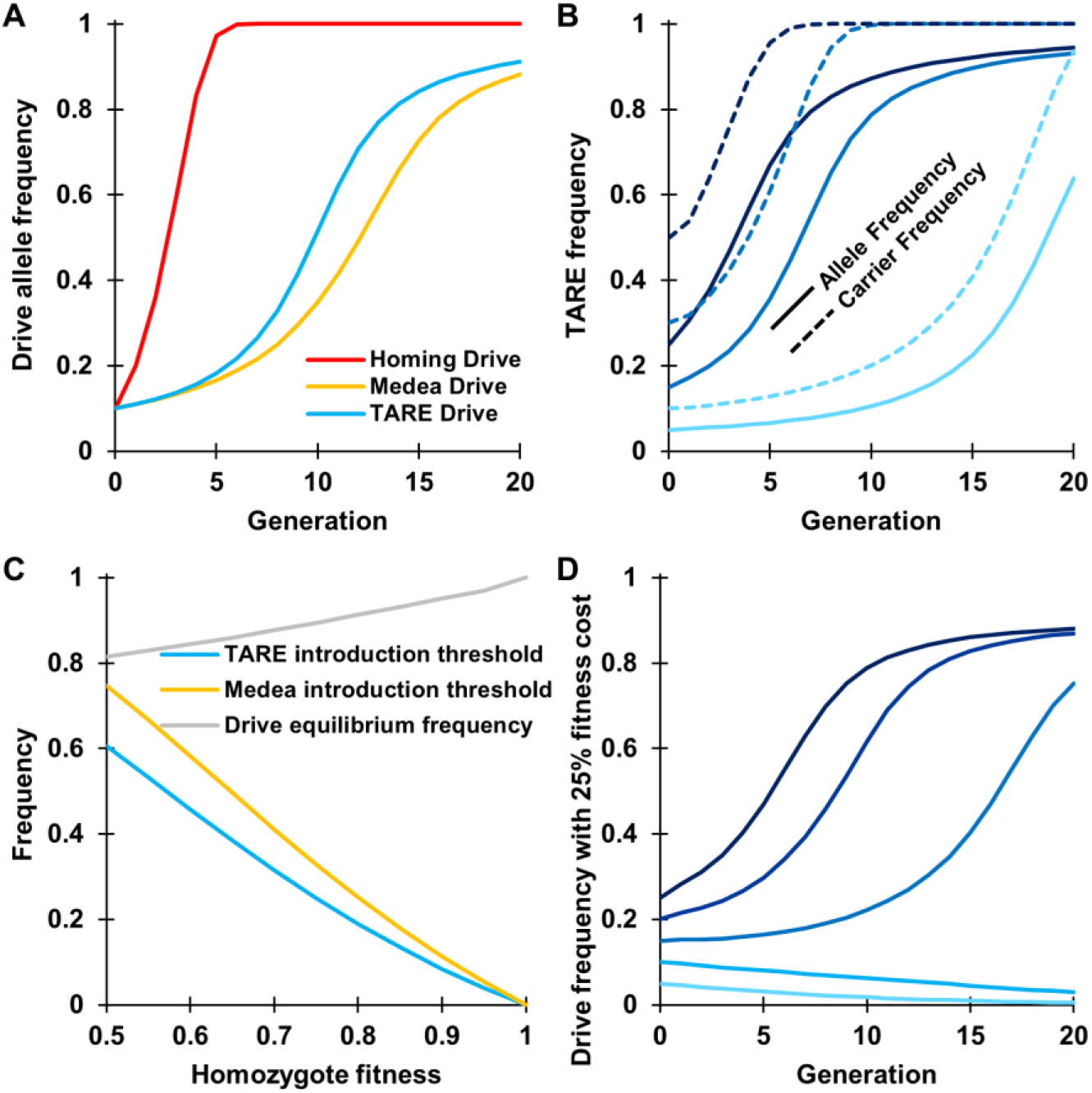
TARE drive dynamics. Expected drive trajectories were simulated in a deterministic model of a single panmictic population with an initial release of drive/wild-type heterozygotes and assuming no fitness costs for ideal drives. (**A**) An ideal TARE drive increases in frequency less rapidly than an ideal homing drive. It has similar dynamics to an ideal *Medea* drive, but with slightly increased speed to fixation since both male and female drive individuals contribute to the disruption and subsequent removal of wild-type alleles. (**B**) A TARE drive is expected to show frequency-dependent dynamics, increasing in frequency more rapidly at moderate frequencies than at low frequencies. At high frequencies, however, the rate at which wild-type individuals are removed is slowed. Nevertheless, the drive should rapidly reach all individuals in a population (in the sense that most individuals should carry at least one copy of the drive) with a moderate initial release size. (**C**) Invasion threshold frequencies of drive heterozygotes as a function of the drive homozygote fitness cost. These thresholds represent unstable equilibria above which the drive increases in frequency and below which the drive is removed. With fitness costs, both TARE and *Medea* drives will not reach fixation, but instead reach an equilibrium frequency as shown, which is the same for both types of drives. Note that all individuals at equilibrium still have at least one copy of the drive allele. (**D**) Drive allele frequency dynamics when assuming a drive homozygote fitness equal to 75% that of wild-type individuals, which yields a threshold heterozygote release frequency of 25% (12.5% allele introduction frequency).

One interesting feature of a TARE drive system is that it should exhibit threshold-dependent dynamics when the drive allele incurs an additional fitness cost to the organism (Figure 2C), similar to the *Medea* system. When introduced above its characteristic frequency threshold, a drive allele is expected to further increase in frequency, whereas it is expected to decrease in frequency and ultimately be lost when introduced below this frequency (Figure 2D). Such threshold-dependent dynamics could be highly desirable for enabling drives to be confined to certain regions, since it would be impossible for the drive to establish in other regions through migration of only a small number of individuals. This is in stark contrast to homing-type drives, which are self-sustaining at any introduction frequency in deterministic models.

Analogous to *Medea*, a TARE drive with any additional fitness costs will also not typically go to fixation in the population but will reach an equilibrium frequency between drive alleles and disrupted alleles (Figure 2C). This is because drive-carrying homozygotes have somewhat lower fitness than drive/wild-type heterozygotes, which is balanced by loss of some offspring without drive alleles when heterozygotes mate with each other. This equilibrium frequency is high unless fitness costs are severe. Even then, all individuals still possess at least one copy of the drive allele once the wild-type has been displaced, which should render TARE drives quite effective for most population modification strategies.

### Drive construct design

We designed a TARE drive at the *h* locus in *D. melanogaster*. Our construct consists of a recoded *h* sequence followed by its natural 3’UTR, a dsRed payload gene driven by the 3xP3 promoter for expression in the eyes (for phenotyping in our *w*^*1118*^ line), and a set of two gRNAs driven by the U6:3 promoter (Figure 3A). The gRNA gene contains a tRNA at the start and another in between the gRNAs, which are spliced out from the RNA transcript, leaving only the mature gRNAs. These gRNAs target a region of *h* downstream from the drive allele (Figure 3B), which prevents copying of the drive allele if gRNA-induced DNA breaks undergo homology-directed repair. Because *h* is a recessive lethal gene, embryos must have at least one functional copy to survive. This can be a wild-type allele, a drive allele, or an r1 resistance allele, in which the *h* gene remains functional despite a change in sequence at both target sites (although we did not detect any such r1 alleles in this study). Any embryo receiving two copies of *h* that have both been disrupted by Cas9/gRNA cleavage will be non-viable. The construct with split Cas9 driven by the germline *nanos* promoter and containing an EGFP reporter was constructed in a previous study (Figure 3C)^17^. This Cas9 allele was located on chromosome 2R, while the drive allele in *h* is on chromosome 3L, so both alleles segregate independently.

**Figure 3.**
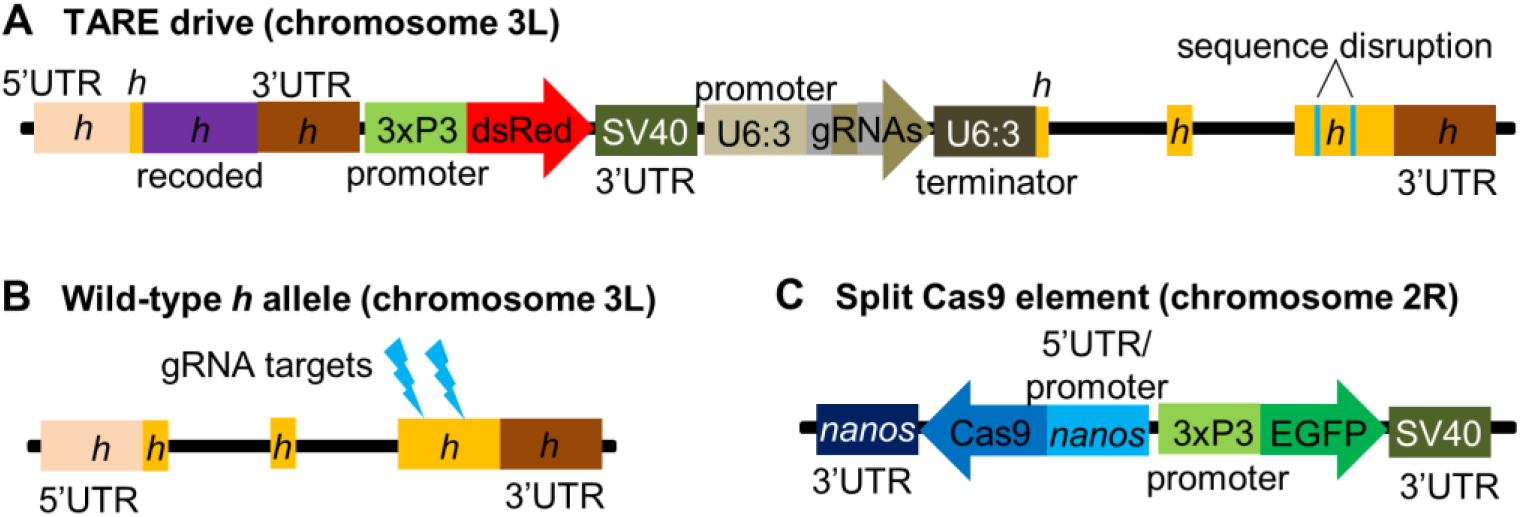
Schematic of the TARE split-drive constructs. (**A**) The split TARE drive is inserted into the coding region of the first exon in *h*. The drive contains a recoded version of *h* and its 3’UTR, a dsRed marker gene driven by a 3xP3 promoter together with a SV40 3’UTR, and a gRNA gene consisting of two gRNAs that target *h*, linked by tRNAs and expressed by the U6:3 promoter. (**B**) The wild-type *h* allele is targeted in the coding sequencing of the third exons by the two gRNAs. (**C**) The supporting element contains Cas9 driven by the *nanos* promoter with a *nanos* 3’UTR, and an EGFP marker gene driven by a 3xP3 promoter together with a SV40 3’UTR.

### Drive evaluation

To assess drive efficiency, we crossed TARE drive homozygote males to females homozygous for the *nanos*-Cas9 allele. The progeny of these were heterozygous for both the drive and the split-Cas9 allele. They were then each crossed to *w*^*1118*^ flies, and the progeny were phenotyped. We found that the progeny of heterozygote females were 87.7% dsRed (Figure 4, Supplemental Data S1), which represented a significant deviation from Mendelian inheritance (p<0.001, Fisher’s exact test). This is likely due to lower viability among flies that did not inherit a drive allele. Nearly all wild-type *h* alleles of these flies were likely disrupted in the germline, and a high proportion of paternal *h* alleles were then disrupted by maternal Cas9 activity, resulting in the death of the embryos where cleavage took place. Embryos that inherited the drive allele would remain viable, regardless of maternal Cas9 activity. These results are further supported by subsequent sequencing of the target locus and analyzing the resulting mosaic sequences (Supplemental Information). We detected wild-type sequences in one out of six flies that inherited the drive allele, but all six sequenced flies that did not receive the drive allele had a detectable wild-type sequence. The progeny of males that were heterozygous for the drive and Cas9 did not show altered inheritance (Figure 4, Supplemental Data S2). Mosaic sequences revealed that only one out of six flies that did not inherit the drive had a fully wild-type target sequence, which supports the notion that most wild-type alleles are cleaved in the germline.

**Figure 4.**
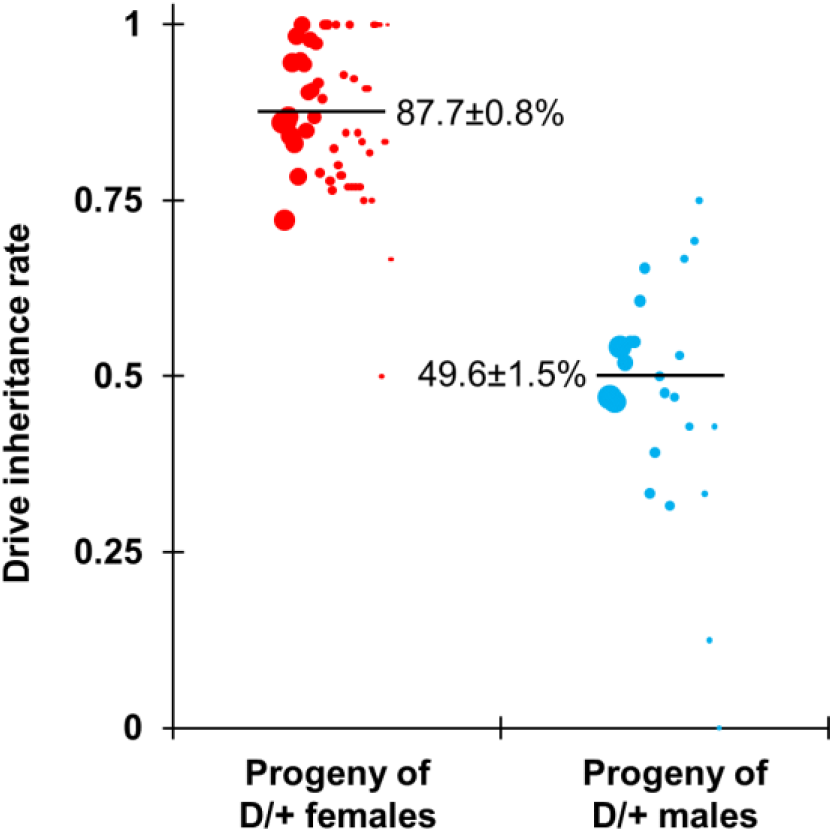
Drive allele inheritance. Females and males heterozygous for the drive and Cas9 were crossed with *w*^*1118*^ individuals, and their progeny were phenotyped for dsRed, which indicates the presence of the drive. Females showed biased inheritance of the drive, since many individuals without the drive had two disrupted copies of *h* and thus were non-viable. Half of the progeny of male drive heterozygotes received the drive, since individuals that received a disrupted copy of *h* still received a functional wild-type copy from their mother that remained undisrupted in the absence of maternal Cas9 activity. The size of the dots represents the sample size of adult progeny from a single drive individual. The horizontal lines specify the average drive transmission.

Flies inheriting the drive from the above cross were most likely heterozygotes for a drive and a disrupted *h* allele. To distinguish germline and maternal Cas9 activity, such heterozygous females that also inherited the Cas9 allele were crossed to *w*^*1118*^ males, and the progeny were then scored as above. In this cross, dsRed inheritance was 95.1%, which was significantly higher than for drive/wild-type heterozygotes (*p*<0.001, Fisher’s exact test). This implies that germline cleavage and disruption is somewhat less effective than 100% for this drive, because the rate of cleavage in the early embryo was likely the same for both crosses.

To confirm the mechanism of action of our TARE drive, we crossed drive/wild-type heterozygotes with one copy of Cas9 with *w*^*1118*^ flies, and also crossed male and female *w*^*1118*^ flies together. Individual flies were then allowed to lay eggs for up to three 20-hour intervals. Eggs were counted at the end of these intervals, and subsequently, pupae were also counted in addition to phenotyping of eclosed adults. Counts of vials adversely affected by fungal growth were discarded to reduce variability in the egg-to-pupae survival rate, though later growing fungus did result in a higher death rate for pupae in some remaining vials when vial fly density was low. We found that female drive heterozygotes had 49.2% egg-to-pupae survival (Figure 5, Data Set S1) and male drive heterozygotes had 79.5% survival (Figure 5, Data Set S2), compared to 82.9% for *w*^*1118*^ individuals without the drive and Cas9 (Figure 5, Data Set S3). Thus, while progeny of male drive heterozygotes had an egg-to-pupae survival rate that was comparable to *w*^*1118*^ individuals, the egg-to-pupae survival rate of progeny from female drive heterozygotes was only about 60% that of *w*^*1118*^ individuals, which was significantly lower (*p*<0.001, Fisher’s exact test) and consistent with our results for drive inheritance in a model in which early embryo Cas9 activity results in the death of most flies not inheriting the drive allele (Data Set S1).

**Figure 5.**
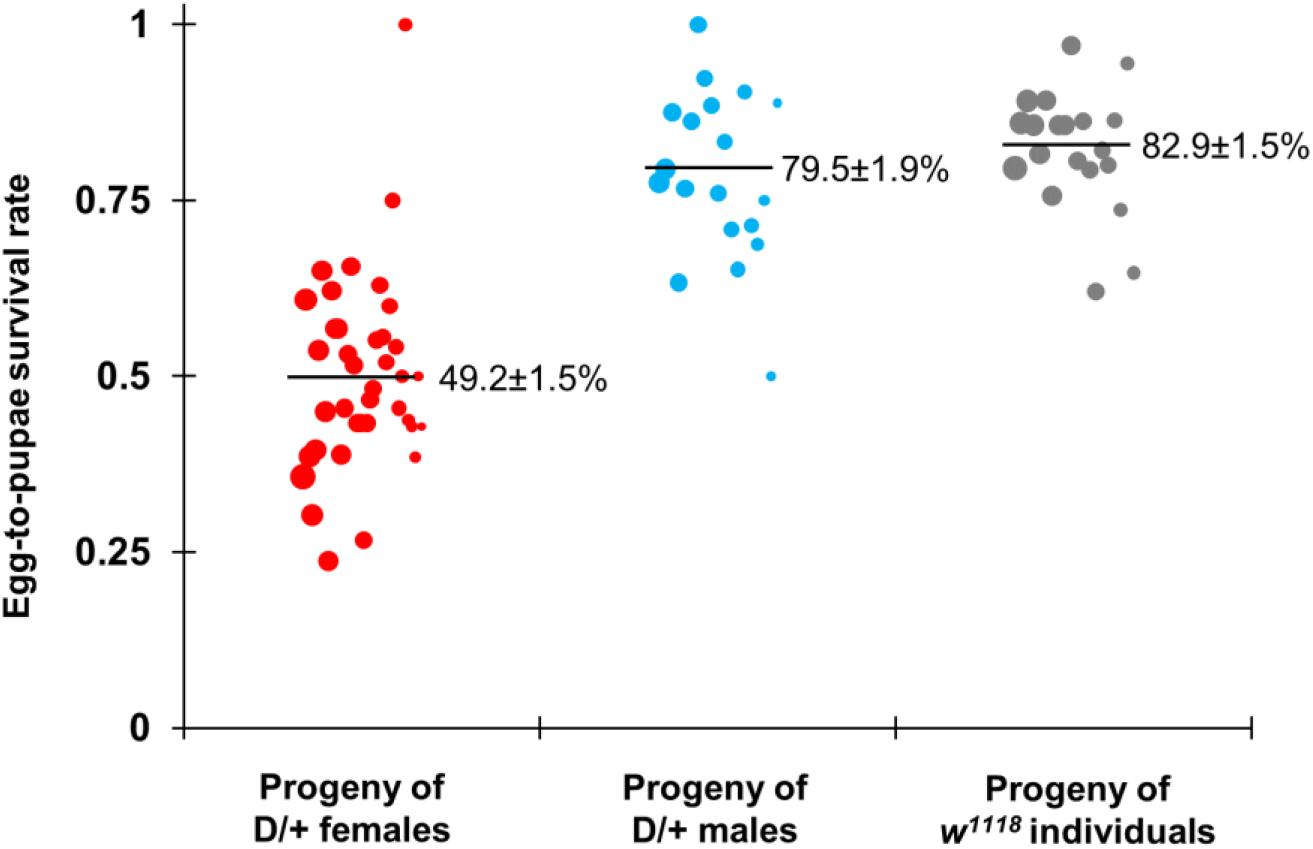
Egg-to-pupae viability. Females and males heterozygous for the drive and Cas9 were crossed with *w*^*1118*^ individuals, and *w*^*1118*^ males and females were crossed together. Eggs were counted in 20-hour intervals, and pupae were counted nine days later. Female drive heterozygotes had substantially lower egg viability, as expected with a mechanism in which maternal Cas9 activity creates recessive disruption in *h*. The size of the dots represents egg sample size from a single female. The average and standard deviation is displayed for each cross.

### TARE drive cage study

To study the performance of the TARE drive in large cage populations, we first crossed drive homozygous males to females homozygous for the Cas9 allele. The progeny were crossed together for several generations, selecting in particular individuals with the brightest dsRed and EGFP phenotype. When individuals were confirmed to be homozygous for both the drive and Cas9 alleles, they were crossed to Cas9 homozygotes with no drive, generating individuals that were drive/wild-type heterozygotes at the drive locus, but still possessing two copies of Cas9. These were crossed to *w*^*1118*^ males, and the resulting dsRed inheritance was 91.1%, which was only slightly higher than for drive/wild-type heterozygotes (*p*=0.027, Fisher’s exact test), most likely because of the increased maternal deposition of Cas9 due to a second copy in the genome. However, this difference was small, implying that the split drive in a Cas9 homozygous genetic background would have similar performance to a complete TARE drive (that includes Cas9) in a wild-type background.

Flies homozygous for both the drive and Cas9 were then expanded and allowed to lay eggs in bottles for one day, and only flies homozygous for the Cas9 allele were allowed to similarly lay eggs for one day in another set of bottles. Flies were then removed, and the bottles were placed in varying proportions in three population cages, which were followed for several generations, with each generation phenotyped for dsRed (Figure 6, Data Set S4). The drive allele reached 100% of the population by the end of the study. Since an r1 allele would be predicted to be paired with an r2 allele approximately 20% of the time (Figure 1), this indicates that either no r1 alleles formed in either cage, or that they were present in very low frequencies (presumably below 0.2%). Both cages outperformed the predictions from an idealized deterministic model assuming perfect drive efficiency and no fitness costs. A caveat to this is that we used a split-drive configuration, and thus, we had no power to detect fitness costs associated with the expression of the large Cas9 protein, which may represent a substantial fraction of a complete TARE drive’s fitness costs. Maternal effects may also have allowed the drive allele to somewhat outperform the predictions of the theoretical model, and the recoded *h* allele itself may be advantageous in a cage setting. None of these factors would likely allow a TARE drive to have a fitness greater than wild-type individuals in a natural setting, particularly if it was carrying a useful payload gene.

**Figure 6.**
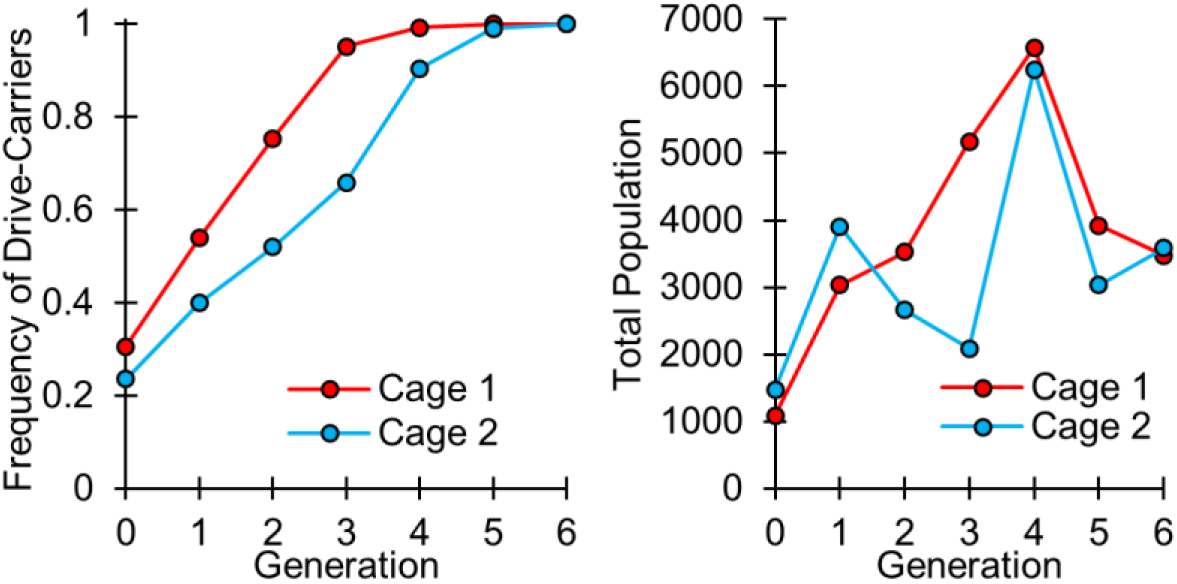
Cage frequency trajectories and population sizes. Individuals homozygous for the drive and Cas9 were mixed with individuals homozygous only for Cas9 in large cages. Cage population were maintained for several discrete generations, and all individuals from each generation were phenotyped for dsRed, indicating that they carried one or two copies of the drive allele.

## DISCUSSION

Our results demonstrate that same-site TARE systems are capable of efficiently modifying large populations without observable resistance evolution. Although we tested our system only in *D. melanogaster*, it should be straightforward to transfer such a system to other organisms, such as mosquitoes, as long as one can find a suitable recessive lethal target gene. In fact, a TARE drive would likely be successful even if lethality or haplosufficiency is incomplete. While we used the promoter of the highly conserved germline-expressed gene *nanos* for Cas9 expression in our construct, a TARE drive should work with a variety of promoters that are active in gametes or their precursor cells, though maternal activity would still be highly desirable for rapid spread. This makes such a system much more flexible than CRISPR homing-type drives, where embryo activity can be problematic due to its propensity for forming undesirable resistance alleles^15,16^. In contrast, such embryo activity actually helps the drive spread faster for a TARE system. As a result, TARE systems are far less prone to the formation of resistance alleles than homing type drives, particularly for population modification drives. We used two gRNAs in our drive construct, but with the same tRNA system, four or more gRNAs could easily be expressed, likely resulting in negligible rates of r1 allele formation with no negative effects on drive efficiency, unlike homing drives^16^.

A TARE drive shows threshold-dependent dynamics, usually with a low threshold. Thus, it would likely spread rapidly in the release region, but fail to establish from occasional long-distance migrants, which could be a useful property of a TARE drive if regional as opposed to global population modification is desired. This is in contrast to homing drives, which in principle, could spread successfully by long-distance dispersal of just a few individuals. Nevertheless, TARE systems are still more invasive than most underdominance systems, which may only persist locally, as opposed to regionally, depending on the type. A common drawback of underdominance systems is that they are typically difficult to construct. By contrast, a TARE-based underdominance system consisting of two TARE alleles, each targeting the gene the other provides rescue for, would presumably be relatively straightforward to design and engineer.

TARE systems do have some limitations, such as the extended period it can take them to go from high frequency to ultimate fixation (or equilibrium). Indeed, a TARE drive will not be predicted to fixate if it has any additional fitness cost, though it would still reach all individuals if there are no r1 resistance alleles. Thus, TARE drive should be particularly suitable for cases in which a single drive allele in all individuals is sufficient to provide the desired population modification effect. This limitation could in principle be overcome by using an X-linked target gene at the cost of slower drive spread^41^. Another limitation of TARE is that it cannot be used for population suppression, though other “TA” systems could possibly do so, at the cost of greater construction difficulty^41^.

TARE systems can be “same-site” or ClvR-type^36^ drives at a “distant-site”. In the former, as demonstrated here, the drive allele is located at the target site gene, while in the latter, the drive allele is located at a different locus in the genome, usually unlinked to the target gene. For the relatively high germline and embryo cut rates demonstrated thus far, both systems have similar population dynamics^41^. However, same-site systems have the advantage that they require an “antidote” element of reduced size, since natural promoter elements are used, enabling easier engineering. This also ensures that a different genomic location and/or an incomplete promoter will not affect expression of the recoded target gene compared to the wild-type gene, enhancing the chance of successful rescue and likely reducing fitness costs. Distant site systems, on the other hand, may be advantageous if the aim is to disrupt a particular gene (other than the target gene), since the gene would be reliably disrupted by the drive allele without the need to target it with additional gRNAs.

Our study shows that TARE systems are promising candidates for regionally-confined population modification drives and do not suffer from the high resistance rates typically observed in homing-type drives. With their great flexibility in choosing promoters and target sites, such drives could potentially be developed in a wide variety of organisms with reduced development time compared to other drive mechanisms. Future studies should investigate TARE drives in other organisms and further explore suitable payloads for potential field deployment.

## ACKNOWLEDGEMENTS

This study was supported by the National Institutes of Health award R21AI130635 to J.C., A.G.C. and P.W.M, National Institutes of Health award F32AI138476 to J.C, and National Institutes of Health award R01GM127418 to P.W.M.

## SUPPLEMENTAL INFORMATION

The following tables show the DNA fragments used for Gibson Assembly of each plasmid and the sequences of DNA oligos. PCR products are shown with the oligonucleotide primer pair used, and plasmid digests are shown with the restriction enzymes used.

| **EGDhg2** | *Template* | *Oligo/Enzyme 1* | *Oligo/Enzyme 2* |
| --- | --- | --- | --- |
| *PCR Product* | pCFD3 | CFD5h_1_F | CFD_1_R |
| *PCR Product* | pCFD5 | CFDh_12_F | CFDh_12_R |
| *PCR Product* | pCFD5 | CFD_2_F | CFD5h_2_R |

| **EGDhg2t** | *Template* | *Oligo/Enzyme 1* | *Oligo/Enzyme 2* |
| --- | --- | --- | --- |
| *PCR Product* | pCFD3 | CFD5ht_1_F | CFD_1_R |
| *PCR Product* | pCFD5 | CFDht_12_F | CFDht_12_R |
| *PCR Product* | pCFD5 | CFD_2_F | CFD5ht_2_R |

| **EGDh2i1** | *Template* | *Oligo/Enzyme 1* | *Oligo/Enzyme 2* |
| --- | --- | --- | --- |
| *PCR Product* | Genomic DNA | hLeft_F | hLeft_R |
| *PCR Product* | gBlock ghr | hcode_F | hcode_R |
| *PCR Product* | Genomic DNA | h3utr_F | h3utr_R |
| *Plasmid Digest* | IHDyi2* | AvrII | XmaI |

\*a plasmid backbone derived from IHDyi2, with the BsiWI site replaced by an XmaI site

| **EGDh2** | *Template* | *Oligo/Enzyme 1* | *Oligo/Enzyme 2* |
| --- | --- | --- | --- |
| *PCR Product* | EGDhg2 | U6_h_F | U6_h_R |
| *PCR Product* | Genomic DNA | hRight_F | hRight_R |
| *Plasmid Digest* | EGDh2i1 | AgeI | XhoI |

### Construction primers

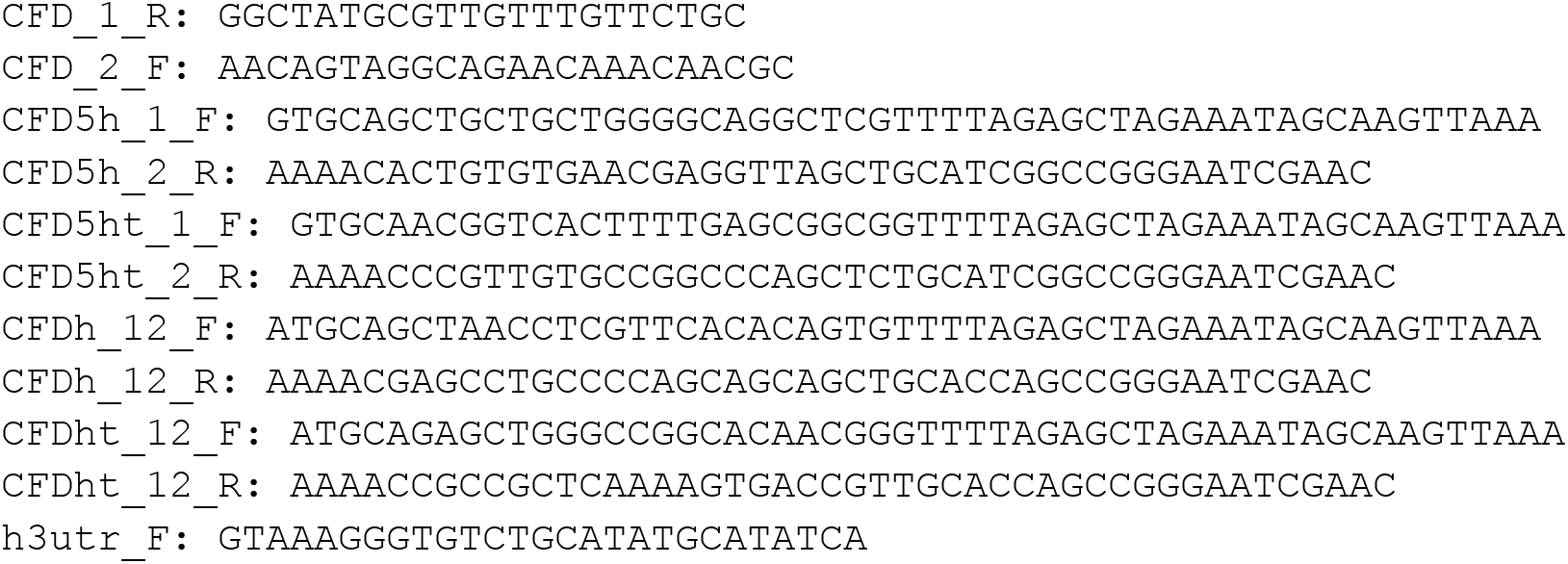
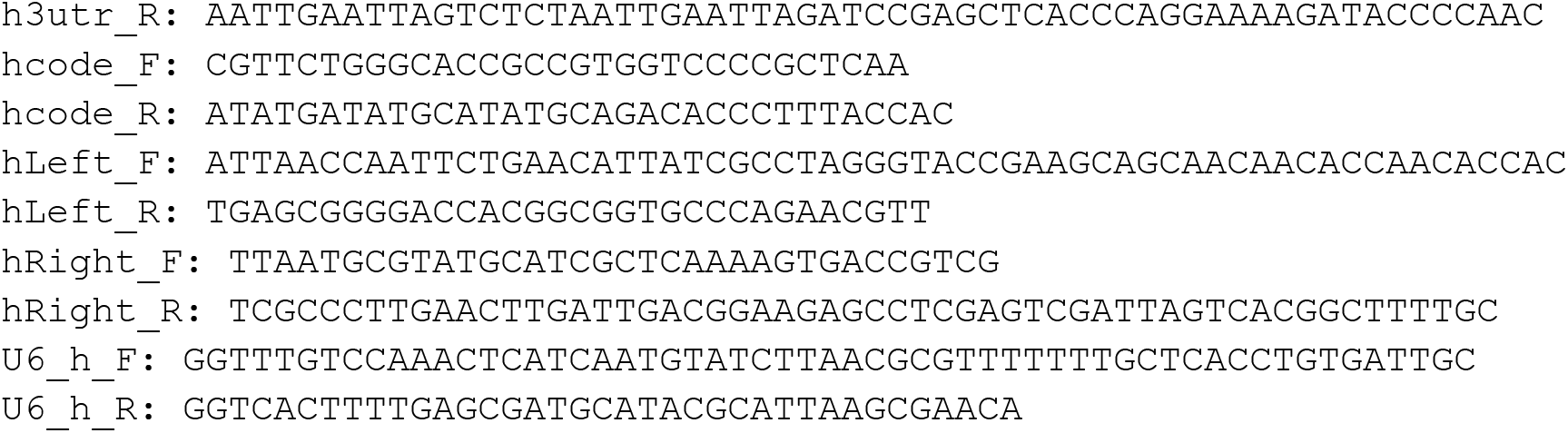

### Sequencing primers

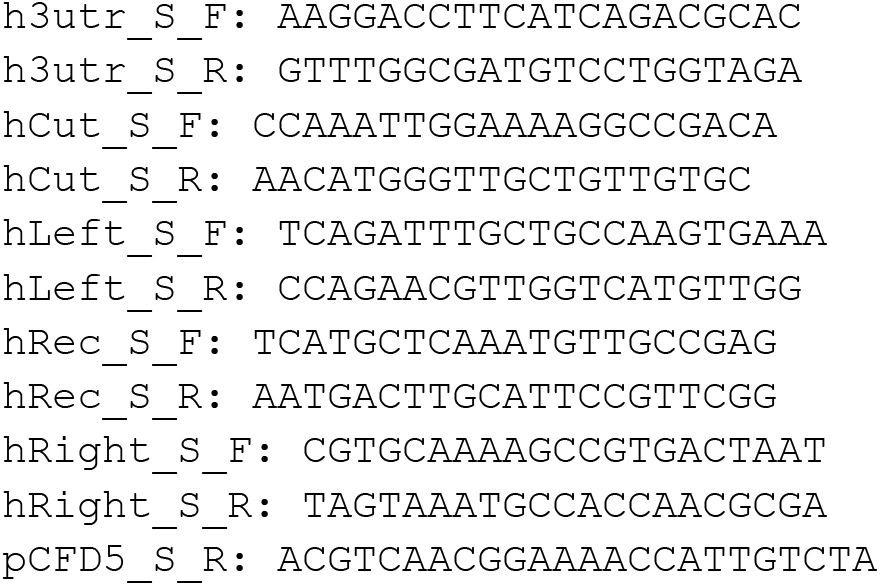

### gBlock

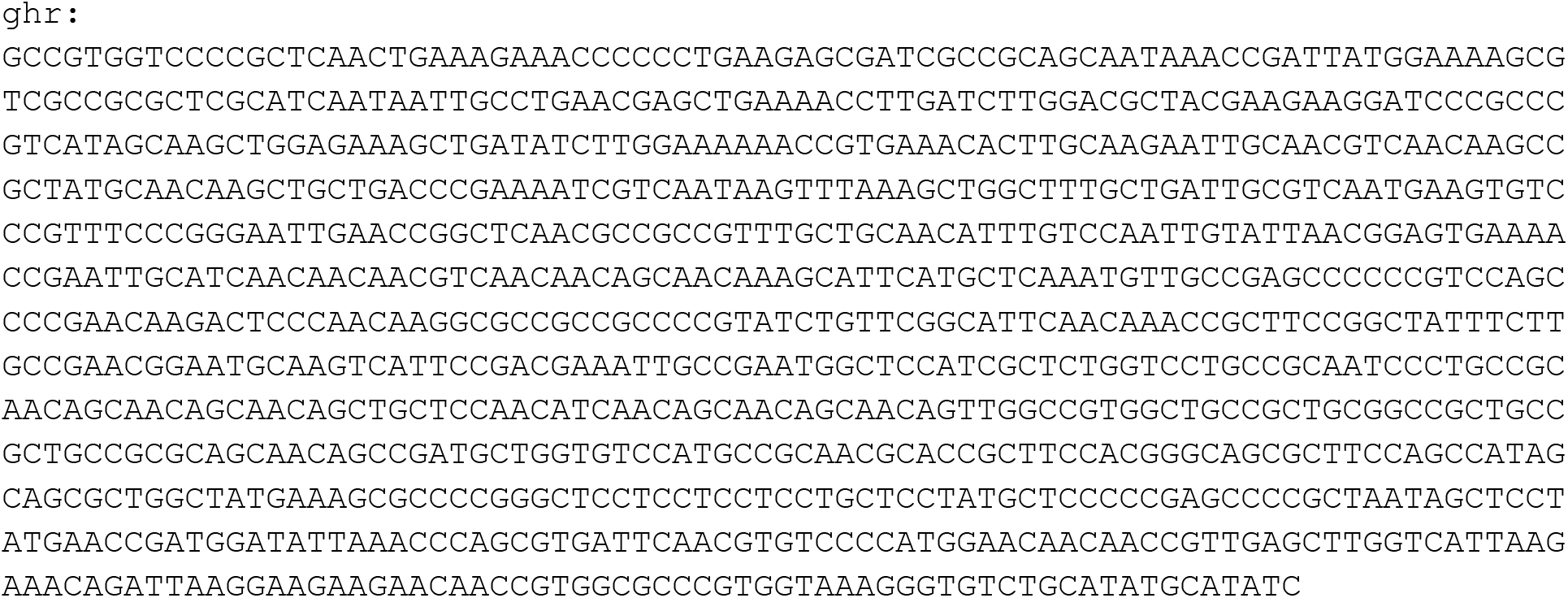

### Sequences of *h* cuts sites for the lines used in the cage experiment

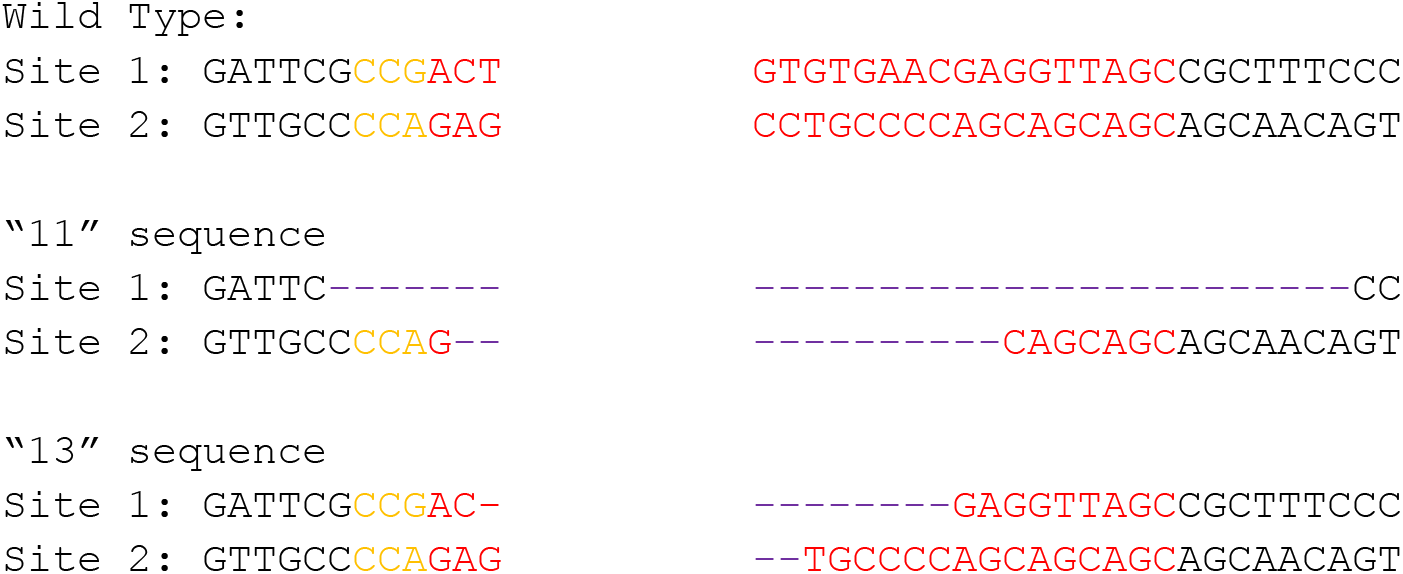

### Other *h* cut site sequence analysis

Due to the presence of multiple sequences, *h* cut sites were distinguished by type (wild-type, mutated, or mosaic). dsRed offspring with a drive mother have a mutated sequence from the drive allele. The table shows the type of the other allele that is present. Similarly, wild-type individuals all had a wild-type allele, and the sequence type shown is the type of other allele.

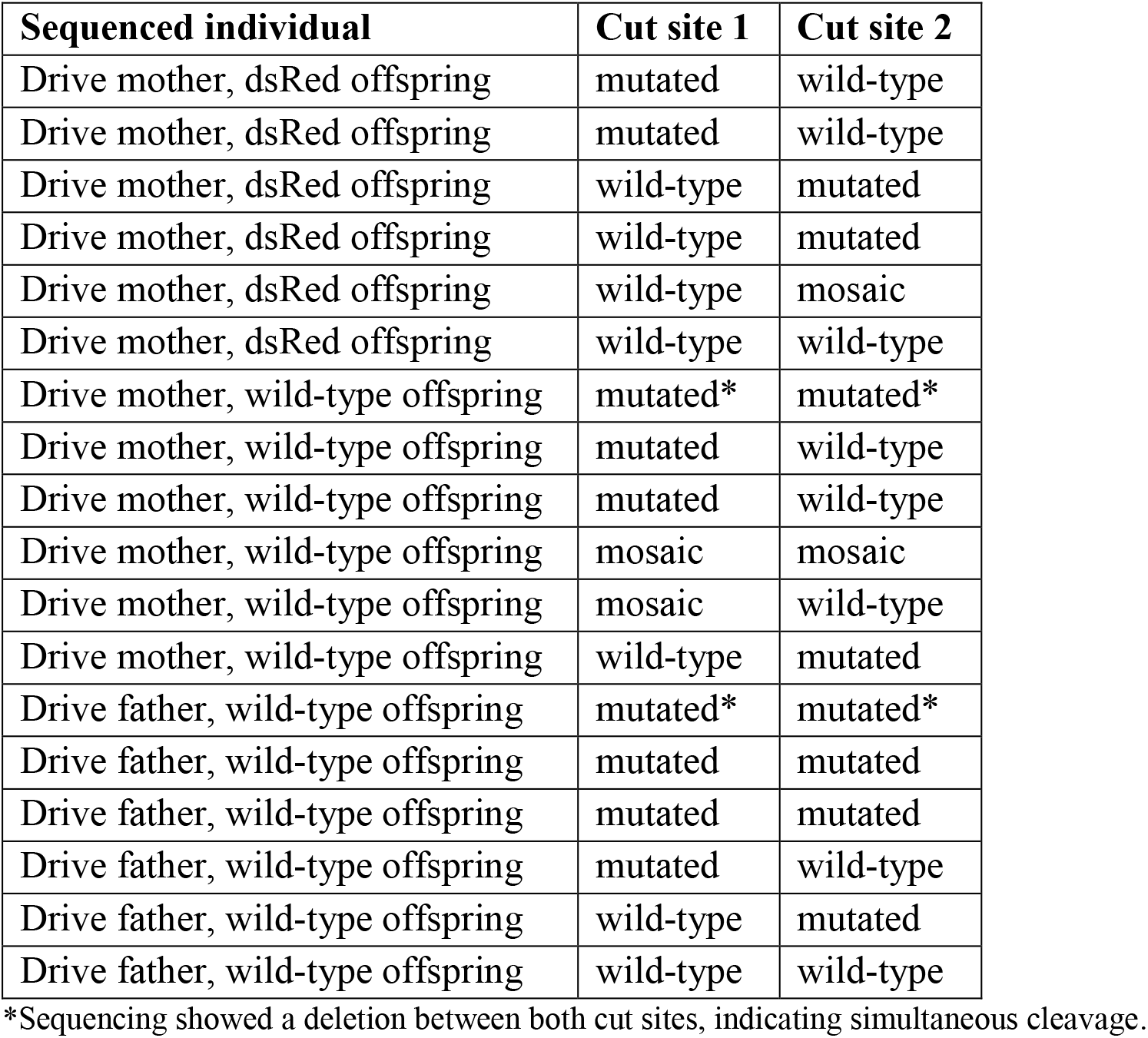

| Sequenced individual | Cut site 1 | Cut site 2 |
| --- | --- | --- |
| Drive mother, dsRed offspring | mutated | wild-type |
| Drive mother, dsRed offspring | mutated | wild-type |
| Drive mother, dsRed offspring | wild-type | mutated |
| Drive mother, dsRed offspring | wild-type | mutated |
| Drive mother, dsRed offspring | wild-type | mosaic |
| Drive mother, dsRed offspring | wild-type | wild-type |
| Drive mother, wild-type offspring | mutated* | mutated* |
| Drive mother, wild-type offspring | mutated | wild-type |
| Drive mother, wild-type offspring | mutated | wild-type |
| Drive mother, wild-type offspring | mosaic | mosaic |
| Drive mother, wild-type offspring | mosaic | wild-type |
| Drive mother, wild-type offspring | wild-type | mutated |
| Drive father, wild-type offspring | mutated* | mutated* |
| Drive father, wild-type offspring | mutated | mutated |
| Drive father, wild-type offspring | mutated | mutated |
| Drive father, wild-type offspring | mutated | wild-type |
| Drive father, wild-type offspring | wild-type | mutated |
| Drive father, wild-type offspring | wild-type | wild-type |

\*Sequencing showed a deletion between both cut sites, indicating simultaneous cleavage.

